# Local protein synthesis is a ubiquitous feature of neuronal pre- and postsynaptic compartments

**DOI:** 10.1101/363184

**Authors:** Anne-Sophie Hafner, Paul G. Donlin-Asp, Beulah Leitch, Etienne Herzog, Erin M. Schuman

## Abstract

There is ample evidence for localized mRNAs and protein synthesis in neuronal dendrites, however, demonstrations of these processes in presynaptic terminals are limited. We used expansion microscopy to resolve pre- and postsynaptic compartments in brain slices. Most presynaptic terminals in the hippocampus and forebrain contained mRNA and ribosomes. We sorted fluorescently labeled synaptosomes from mouse brain and then sequenced hundreds of mRNA species present within excitatory boutons. After brief metabolic labeling, more them 30% of all presynaptic terminals exhibited a signal, providing evidence for ongoing protein synthesis. We tested different classic plasticity paradigms and observed unique patterns of rapid pre- and/or postsynaptic translation. Thus presynaptic terminals are translationally competent and local protein synthesis is differentially recruited to drive compartment-specific phenotypes that underlie different forms of plasticity.

**One sentence summary:** Protein synthesis occurs in all synaptic compartments, including excitatory and inhibitory axon terminals.

---

Neurons are morphologically complex possessing a typical cell body from which emerges elaborately branching dendrites and axons. Indeed, most of a neuron’s area is accounted for by its dendrites and axons: for example, the dendrites of rodent pyramidal neuron exceed 10 mm in length (*1*) and a human axon (e.g. in the sciatic nerve) can reach up to 1 meter length. The massive network represented by dendrites and axons provides the surface area to accommodate the 1000-10000 synapses, both excitatory and inhibitory, typically formed by an individual neuron. At synapses, the complement of proteins present represents the best phenotypic indicator of both the type and strength of the synapse. The regulation of synaptic proteins, by post-translational modifications and by ongoing protein synthesis and degradation, drives homeostasis and plasticity at synapses (*2–4*).

It has been proposed that a substantial fraction of proteome supply and remodeling occurs locally within synapses (*5–8*). While a wealth of data has led to consensus that protein synthesis occurs in mature dendrites (*7*, *9*) there has been much less agreement about local translation in mature axons. Many studies have shown that local translation is required for axonal growth during development and repair following injury (e.g. (*10–14*)). In addition, a few recent studies have shown that mature retinal ganglion cell axons contain competent translational machinery and mRNAs (*15*) or use presynaptic translation during plasticity at certain synapse types (*16*, *17*). In addition, there is ample evidence that invertebrate axons can synthesize proteins (e.g. (*18*)). Despite these data, controversy has arisen from an inability to detect reliably ribosomes in axons or terminals (*19*, *20*) and the persistent idea that axonal protein needs are adequately served by the well-documented system of axonal transport (e.g. (*21*)).

Here, to determine whether translation in axon terminals is a common feature of mature brains, we used advanced microscopy methods to determine the abundance and diversity of the components required for translation in nerve terminals from multiple mouse brain areas. We also purified a molecularly-defined population of mature presynaptic nerve terminals and directly sequenced the resident mRNA population. Finally, we conducted high-resolution metabolic labeling to ascertain the frequency of protein synthesis events in all synaptic compartments.

## Presynaptic terminals from mouse cortex and hippocampus contain translation machinery

Efforts to localize molecules or cell biological events to neuronal pre- or postsynaptic compartments using fluorescence microscopy are limited by the tight association of the axonal bouton and the dendritic spine or synapse; the synaptic cleft, the only clear region of separation, is only ~ 20 nm wide. Here, in order to increase the resolving power to visualize mRNA molecules in pre- and postsynaptic compartments, we optimized fluorescence in situ hybridization (FISH) and nascent protein detection methods for use with expansion microscopy (*22*) (Fig. 1A; see Methods). We used adult mouse brain slices or rat cultured hippocampal neurons (DIV 18-21) and found that expansion resulted in an enlargement of both pre- and postsynaptic compartments, with an average expansion of ~3.4 fold. This yielded a clear separation between the pre- and postsynaptic compartments. To evaluate whether ribosomes and mRNA species are present in defined presynaptic compartments, we used immunolabelling for either excitatory (vGLUT1; (*23*, *24*) or inhibitory (vGAT; (*25*, *26*)) nerve terminals in expanded mouse brain sections (both cortex and hippocampus) (Fig. 1B-E) or rat cultured hippocampal neurons. We took care to identify the molecules-of-interest within individual z-sections positively labeled for excitatory or inhibitory terminals. We noted that signal detected outside of immunolabeled compartments corresponded to signal arising from nearby unlabeled cells. We detected ribosomes in a large majority (>75%) of both excitatory and inhibitory presynaptic nerve terminals, using antibodies directed against either a small (RPS11) or a large (RPL26) ribosomal protein (Fig. 1 B-E). Next, we used FISH probes to detect 18s and 28s rRNA as well as polyadenylated mRNA (detected with a poly d(T) probe) in expanded samples (Fig. 1B-E). Consistent with the abundance of ribosomal proteins, we detected rRNA in over 80% of both excitatory and inhibitory nerve terminals (Fig. 1B-E). RNase treatment effectively reduced all rRNA signal. In cultured neurons, we also noted that poly(A) mRNA was abundant, as expected, in dendritic spines. In addition, we used an anti-tau antibody to label axons and detected both 18s and 28s rRNA in axonal segments. Thus, mRNAs and ribosomes were abundant in excitatory and inhibitory presynaptic nerve terminals from both mouse brain slices and rat hippocampal cultured neurons.

**Fig. 1.**
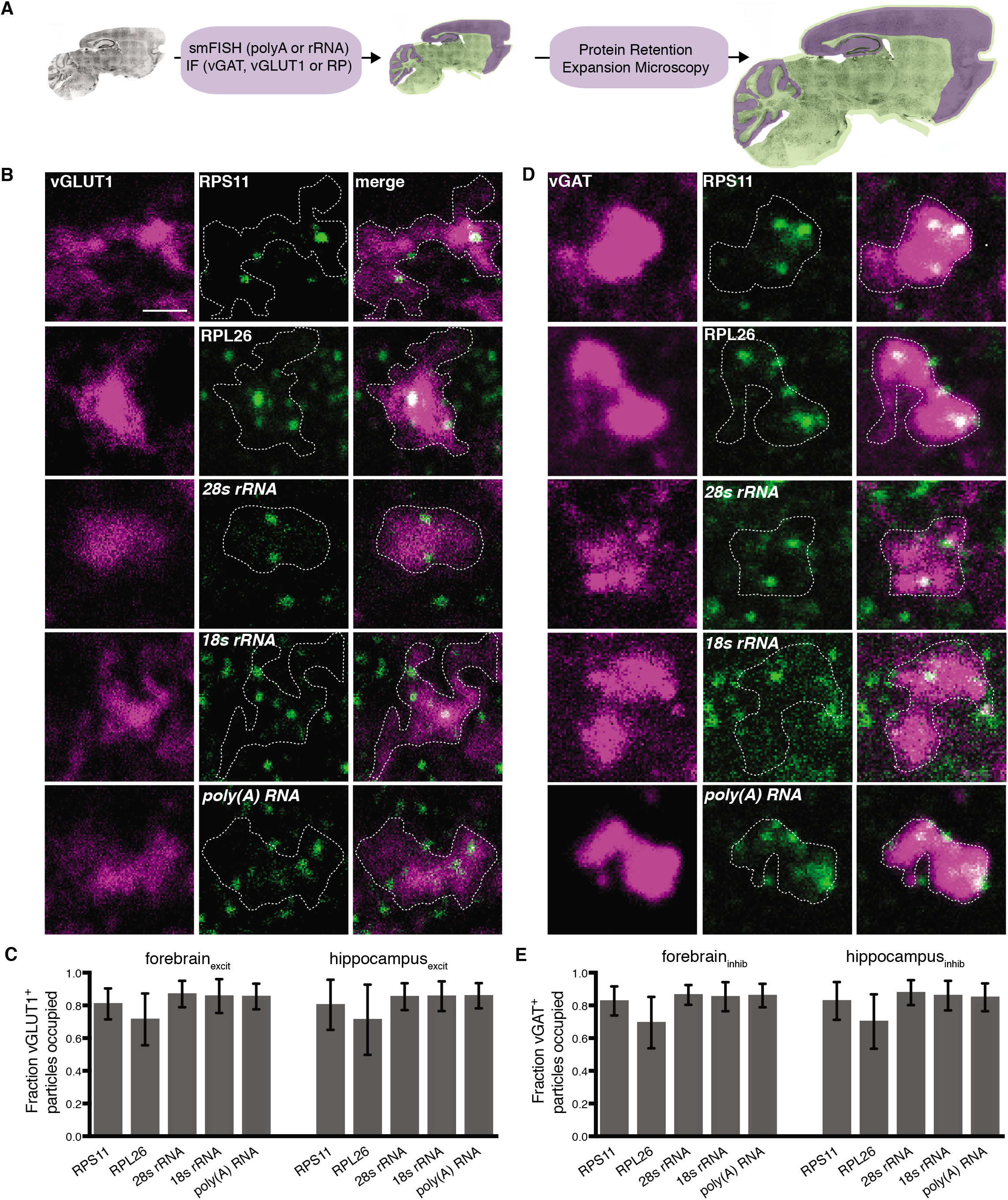
Abundant ribosomes and mRNA in presynaptic compartments of the mature mouse forebrain and hippocampus. (**A**) Scheme indicating the experimental workflow. Sagittal brain slices from adult mice were processed for FISH and fluorescent immunostaining (IF) for presynaptic terminal types, excitatory (vGLUT1) or inhibitory, (vGAT) and then subjected to a protein retention expansion microscopy protocol (see Methods). For illustration purposes, a brain section is pseudo-colored to indicate the distribution of RNA (lavender) and vGLUT1 (green). (**B**) Representative images of expanded presynaptic compartments chosen for their positive vGLUT1^+^ or (**C**) vGAT^+^ signal (magenta) showing the presence of both small and large ribosomal proteins (green; in first trio of images), large and small ribosomal RNA (green; in middle trio of images) or poly(A) RNA (green; in last trio of images) as well as the merged images showing both signals (last image in each trio). Scale bar = 1.5 um. (**D**) Bar graph showing analysis for all vGLUT1^+^ compartments analyzed from forebrain and the hippocampus. Over 75% of all vGLUT1^+^ terminals contained ribosomal proteins and RNA as well as poly (A) RNA. Data acquired from 4 different animals per condition, mean and SEM are plotted. (**E**) Bar graph showing analysis for all vGAT^+^ compartments analyzed from forebrain and the hippocampus. Over 75% of all vGAT^+^ terminals contained ribosomal proteins and RNA as well as poly (A) RNA. Data acquired from 4 different animals per condition, mean and SEM are plotted

## Isolation and characterization of vGLUT1+ terminals from the adult mouse brain

The presence of poly(A) mRNA in axon terminals suggested the capacity for protein synthesis. However, it did not indicate the breadth of translational machinery or the mRNA population potentially available for translation in identified synapse types. In order to characterize transcripts and translational machinery in excitatory presynaptic terminals, we used our recently developed platform that couples fluorescence-sorting with biochemical fractionation to sort and purify fluorescently labeled synaptosomes (fluorescence-activated synaptosome sorting; FASS) (Fig. 2A). The “pre-sorted” and sorted synaptosomes comprise resealed presynaptic synaptic compartments, sometimes associated with an “open” postsynaptic membrane (*27–29*) (Fig. 2B); the sorted synaptosomes also lacked dendritic and ER elements. Starting with the forebrain of adult vGLUT1^venus^ knock-in mice, in which all vGLUT1^+^ synapses were fluorescently labeled (*30*), we prepared and sorted vGLUT1^+^ synaptosomes for FISH, immunocytochemistry (Fig. 2C-I) and ultimately RNA sequencing (Fig. 3). We first examined whether the vGLUT1^+^ sorted synaptosome population, reflecting the composition of excitatory synapses in vivo, possessed the molecular elements that we detected in the expanded hippocampal and forebrain tissue (Fig. 1B-C). Using sparse plating of individual vGLUT1^+^ synaptosomes combined with imaging we determined the incidence of poly(A) mRNA and ribosomal proteins together with a postsynaptic density marker protein, PSD-95 (Fig. 2C-I). Over 80% of all sorted vGLUT1^+^ synaptosomes contained poly(A) mRNA (Fig. 2C,E), ribosomal proteins (Fig. 2D,G,I) and rRNA. As expected, a smaller fraction (~60%) of synaptosomes were associated with PSD-95 (Fig. 2C,D,G). To examine whether this is a universal feature of excitatory synapses we also examined ribosomal protein labeling in vGLUT1^+^ synaptosomes sorted from the adult mouse hippocampus or cerebellum. We found a strikingly similar high occupancy of ribosomes in the hippocampal and cerebellar excitatory synapses: ~90% were positive for ribosome immunolabeling (Fig. 2I). In addition, as observed in adult brain slices, a large majority of the vGAT^+^-immunolabeled synaptosomes were also immunopositive for ribosomes.

**Fig. 2.**
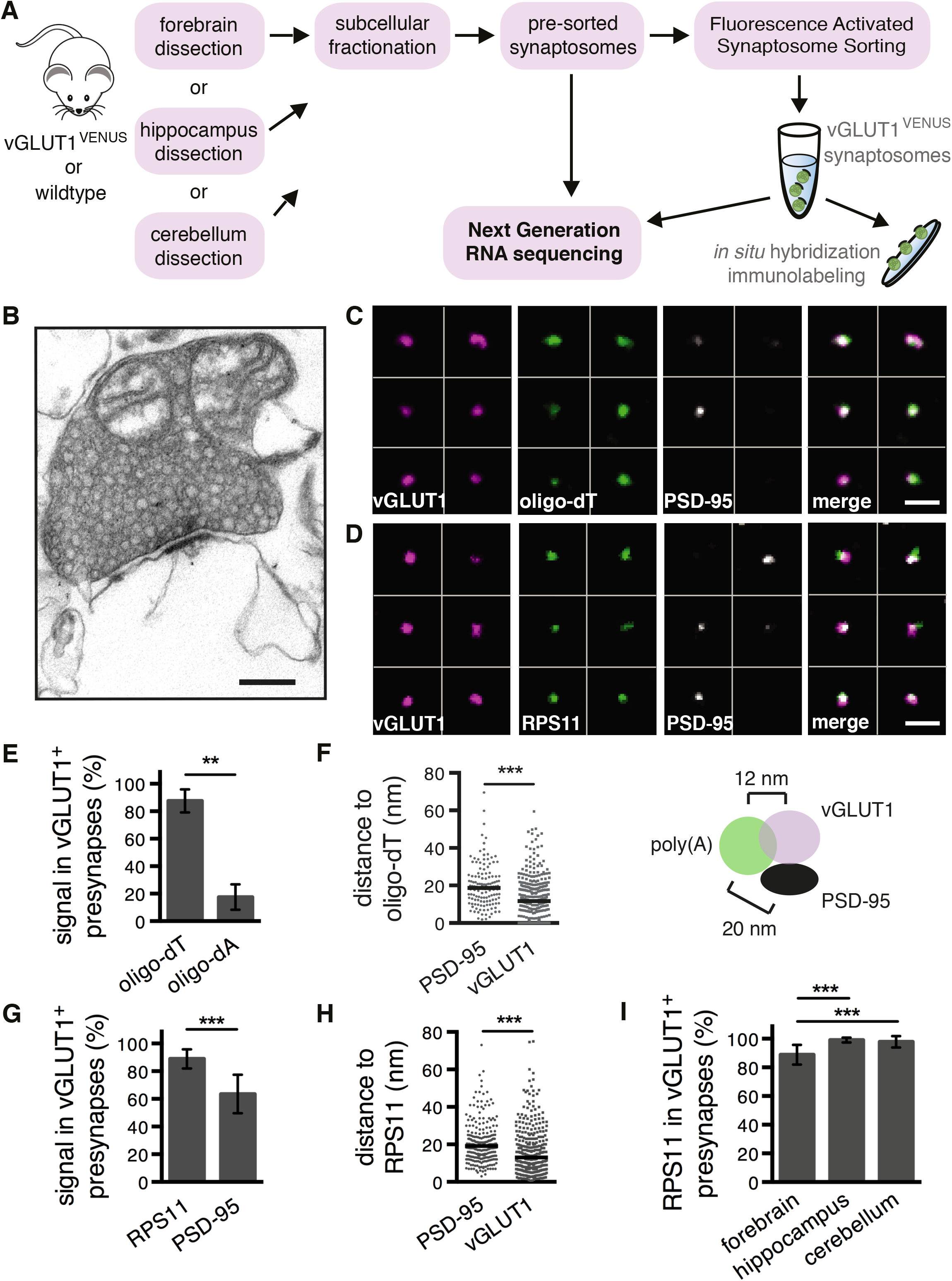
Presynaptic compartments isolated from adult mouse brain contain spatially organized mRNA and ribosomes. (**A**) Scheme showing experimental flow for analysis of the global (pre-sorted) synaptosome and fluorescently sorted vGLUT1^+^ synaptosome population. (**B**) Representative electron micrograph of a pre-sorted synaptosome. Scale bar = 200 nm. (C-D) Example confocal images of sparsely plated vGLUT1^+^ synaptosomes labeled by FISH and immunofluorescence showing the vGLUT1^+^ signal in each synaptosome panel as well as the presence or absence of poly(A) (**C**) or RPS11 (**D**) and PSD-95 for the same samples. Merged images are shown in the last panel. Scale bar = 5 μm. (**E**) Bar graph showing analysis for all synaptosomes analyzed (analysis includes sorted and pre-sorted as both populations yield similar results) (n^dT^ = 921 and n^dA^ = 1069 from 2 biological replicates) showing that 87.6% +/− 8.4 of all VGLUT1^+^ terminals were positive for a FISH oligo-dT probe while less than 17.5% +/−9.2 were positive for a FISH oligo-dA probe. ** Unpaired t-test, p ≤ 0.01. (**F**) Plot of all data points for sorted synaptosomes and median (left) and a scheme (right) showing center-to-center distance between fluorescent signals. The center of the oligo-dT signal (green) was on average 12.8 nm from the center of the vGLUT1 signal (lavender) and 20.0 nm from the PSD-95 signal (black). (n^vGLUT1^ = 317 and n^PSD-95^ = 134) *** Unpaired t-test, p ≤ 0.001. (**G**) Bar graph showing analysis for all synaptosomes analyzed (analysis includes sorted and pre-sorted) (n = 568 from 3 biological replicate experiments) showing that 88.9% +/−6.9 of all vGLUT1^+^ terminals were positive for RPS11 and 63.5% +/−14.0 were positive for PSD-95. *** Unpaired t-test, p ≤ 0.001. (**H**) Plot showing the center-to-center distances between fluorescence signals. The center of the RPS11 signal was on average 19.5 nm from the PSD-95 signal and 15.9 nm from the center of the vGLUT1 signal. *** Unpaired t-test, p ≤ 0.001. (**I**) Bar graph showing analysis for all synaptosomes analyzed (forebrain data are the same as in 2F, for hippocampus and cerebellum we analyzed vGLUT1^+^ sorted synaptosomes) (n^forebrain^ = 568, n^hippocampus^ = 834 and n^cerebellum^ = 236 from at least 2 biological replicate experiments) showing that greater than 80% of all vGLUT1^+^ terminals were positive for RPS11 in all 3 brain regions (forebrain = 88.9% +/− 6.9; hippocampus = 99.1% +/− 1.4; cerebellum = 97.9% +/− 4.0). *** Kruskal-Wallis nonparametric test followed by Dunn’s multiple comparison test, p ≤ 0.001. All data are shown as mean +/− SD.

We took advantage of the punctate nature of the imaged fluorescent signals to calculate the center-to-center distances for poly(A) mRNA or RPS11 and vGLUT1 or PSD-95 proteins. Using stimulated emission depletion microscopy (STED) we confirmed the tight spatial relationship between vGLUT1 and RPS11. The measured distances were consistent with the localization of the presynaptic translation machinery as slightly offset from the synaptic cleft (Fig. 2H-I). This suggests that presynaptic translation occurs away from the active zone. In addition, we optimized expansion microscopy for application to synaptosomes and further probed the spatial organization of the ribosome population relative to the active zone. Synaptosomes were visualized using markers differentially localized within boutons: vGLUT1 (synaptic vesicle pool), bassoon (soluble scaffolding protein) and RIM1 (active zone membrane protein). Consistent with the above data, in the expanded synaptosomes ribosomes (measured with 18s and 28s FISH) were positioned closer to vGLUT1 and Bassoon than the active zone membrane (measured with RIM1 immunolabeling) (Fig. 3A-F). Finally, we used immunoEM to detect the ribosomal protein RPS11 in the pre-sorted synaptosome. We detected RPS11 in a majority of the presynaptic terminals (Fig. 3G-J) and the localization, again offset from the active zone, is consistent with the above data. Thus, the majority of presynaptic terminals, both excitatory and inhibitory, contained both poly(A) mRNA and ribosomal protein, indicating a clear capacity for protein synthesis.

**Fig. 3.**
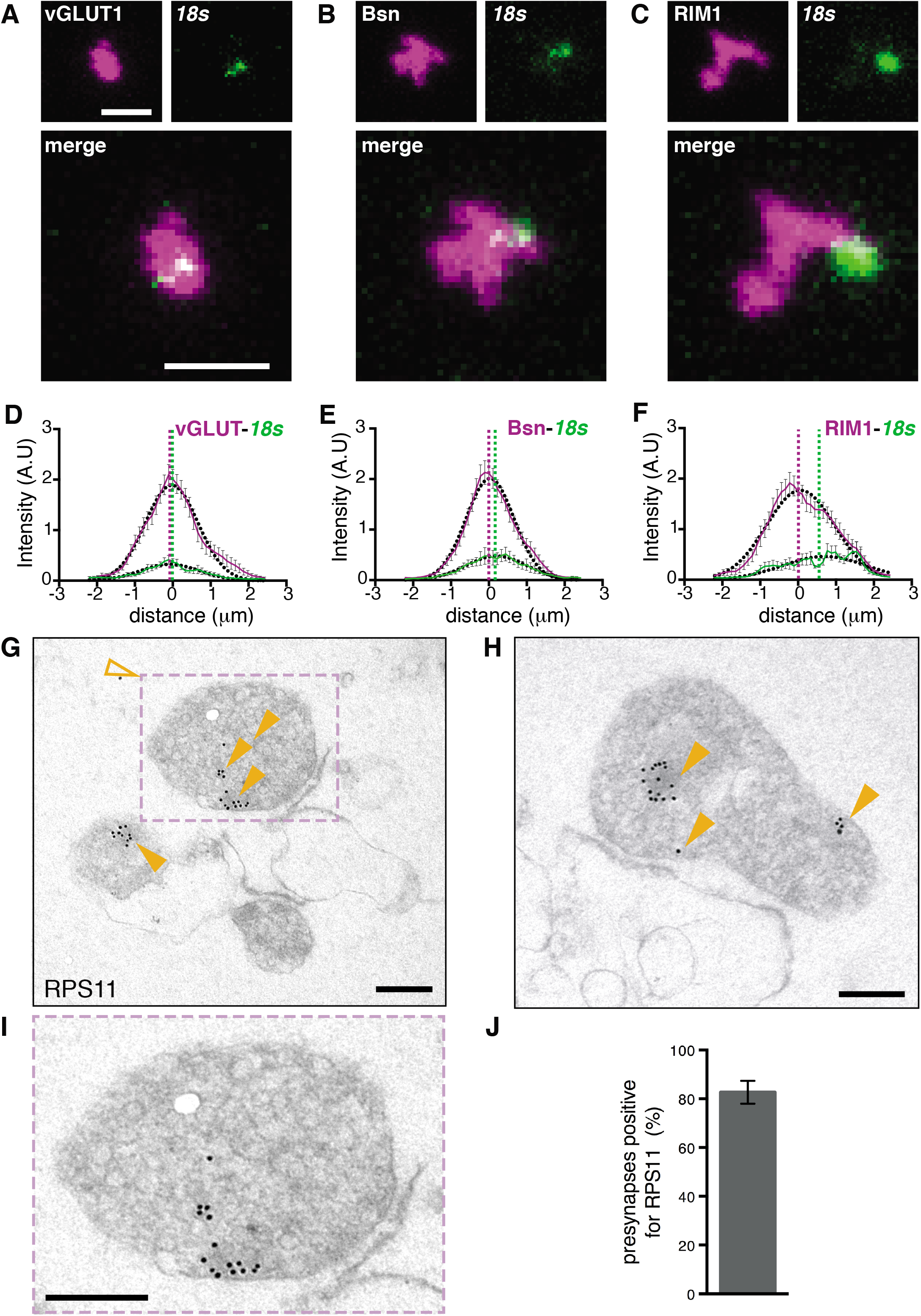
Ribosomes are spatially offset from the active zone in presynaptic terminals. (**A-C**) Synaptosomes were immunostained for vGLUT1 (**A**), bassoon (**B**) or RIM1 (**C**) (magenta) and processed for FISH against 18s rRNA (green). Scale bar = 2.5 um. (**D-F**) Line scan analysis was performed (see methods) to assess the relative spatial distribution of ribosomal RNA in the synaptosomes to the pool of synaptic vesicles or the active zone. Graphs depict the signal distribution of either the presynaptic marker (solid magenta curve) or the ribosomal RNA (solid green curve). Dotted lines indicate the peak of the Gaussian fits (black dotted curves), indicating that the ribosomes were offset from the active zone. n= 40 synaptosomes analyzed from 3 biological replicates. Error bars correspond to SEM. Scale bar = 2.5 um. (**G-I**) Representative electron micrographs of pre-sorted synaptosomes after post-embedding immunostaining using an anti-ribosomal protein RPS11 antibody. (**G**) Electron micrograph showing 3 terminals, 2 of which were associated with a postsynaptic membrane, and 2 of which contained multiple gold particles reflecting the presence of ribosomes in these compartments (solid arrows). Note that there was only 1 gold particle in the field of view outside of terminals (open arrow). Scale bar = 200 nm. (**H**) Electron micrograph showing a terminal containing multiple gold particles (solid arrows) and an open postsynaptic compartment. Scale bar = 200 nm. (**I**) Enlarged view of the terminal in the boxed area in (**G**), suggesting that ribosomes are not present in the active zone. Scale bar = 200 nm. (**J**) Bar graph showing RPS11 occupancy in synapses from immunoEM images (82.7 +/− 4.7, n = 2 biological replicates and 147 terminal counted). Data are shown as mean +/− SD.

## The presynaptic transcriptome of vGLUT1+ terminals

To discover the transcriptome present in adult mouse presynaptic boutons, we used RNA sequencing to identify the mRNA population of both the pre-sorted and vGLUT1^+^ synaptosomes (see Methods). From 3 biological replicates for each group (pre-sorted and sorted), we obtained in total 244 Mio reads that, following genome alignment, yielded 12,730 transcripts detected in all replicates from both groups (196 Mio uniquely mapped reads in total) (Fig. 4A). We analyzed the transcripts that were significantly enriched or depleted in the vGLUT1^+^ sorted population (relative to the pre-sorted synaptosomes) and identified 468 and 792 transcripts, respectively (Fig. 4A-B). Enriched transcripts overlapped to varying degrees with prior synaptic sequencing studies. Gene ontology analysis of the vGLUT1^+^ enriched transcripts (using the ~12.7 K transcripts from the input forebrain transcriptome) revealed a significant overrepresentation of genes coding for presynaptic active zone proteins, ribosomal proteins and other groups like synapse (Fig. 4C). Amongst the most enriched in the vGLUT1^+^ presynaptic transcriptome were many well-known presynaptic proteins including *Bassoon* (*Bsn*), *Rims1-3, Stx6*, as well as signaling molecules like *Sergef* and *Rapgef4* (Fig. 4B,F) and mitochondrial proteins. Amongst the 792 transcripts depleted in the vGLUT1^+^ transcriptome were many coding for neurotransmitter receptors of the GABA and AMPA family, indicating the depletion of postsynaptic and dendritic components through our synaptosome sorting (Fig. 4D,F). Also, transcripts coding for membrane proteins including endoplasmic reticulum proteins, such as *Ergic1, Calr* or *Sec62*, were diminished in the vGLUT1^+^ sorted synaptosomes. There was also a clear depletion of transcripts coding for integral synaptic vesicle proteins (Fig 4D,F), consistent with a recent report of somatic vesicle biogenesis and transport (*31*). Of note, among the 468 transcripts enriched in vGLUT1^+^ terminals, 62 were known targets of the RNA-binding Fragile X Mental Retardation Protein (FMRP), the loss of which causes fragile X syndrome (Fig. 4E) (*32*). We validated the presence (or absence) of several of the vGLUT1^+^ enriched transcripts including *Rapgef4, Adcy1, Bsn, Kif5a*, or *Actb* in sparsely plated vGLUT1^+^ synaptosomes using FISH (Fig. 4G,H). Thus presynaptic compartments from adult mouse forebrain contained the requisite machinery and a diverse mRNA population for protein synthesis.

**Fig. 4.**
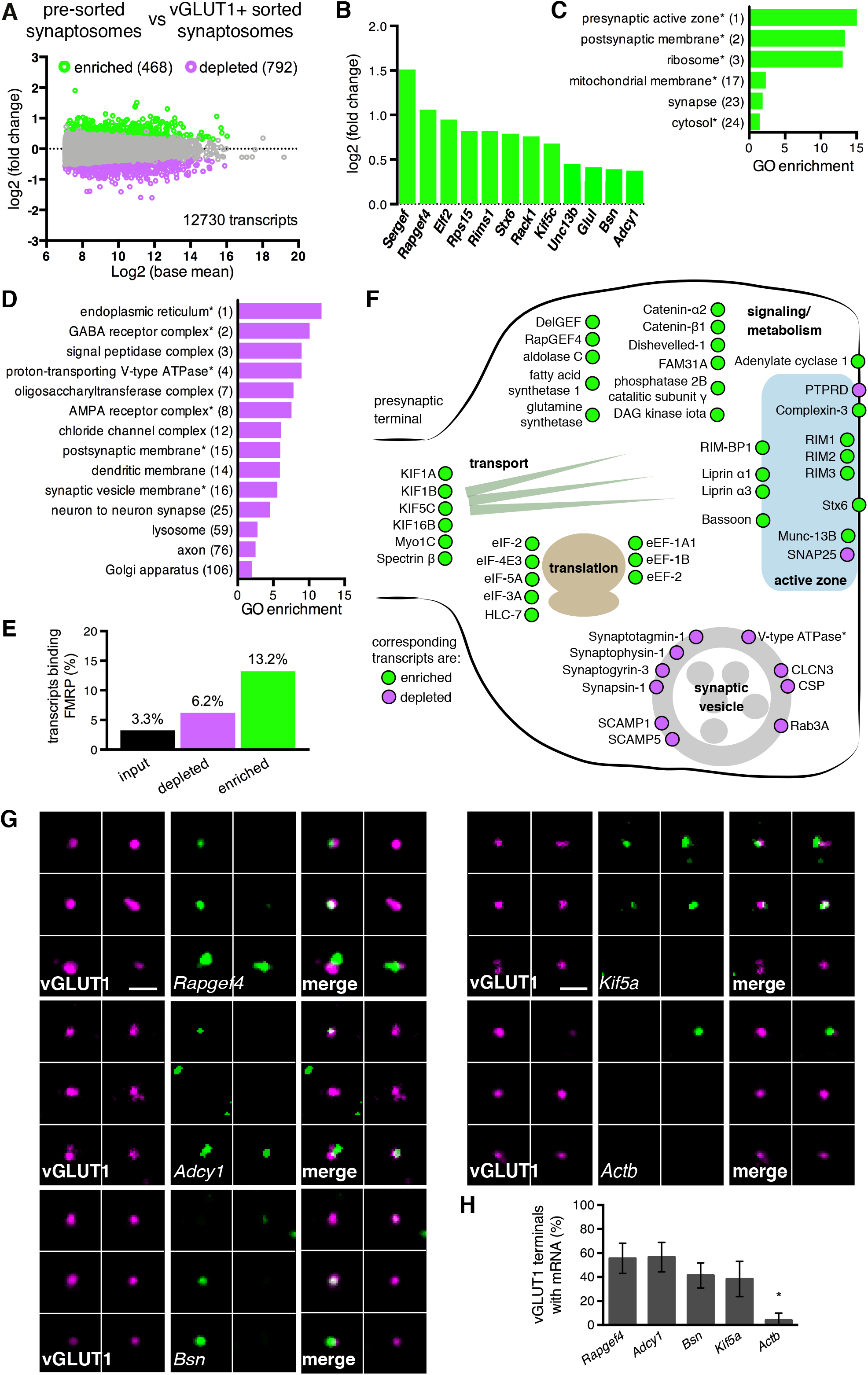
An excitatory presynaptic transcriptome from mature mouse synapses. (**A**) Differential expression analysis (see Methods) showing the relationship between expression (reads per million) and the significant enrichment (green dots) or depletion (magenta dots) in the vGLUT1^+^ sorted vs. pre-sorted synaptosomes. (**B**) Selected list of significantly enriched individual vGLUT1^+^ presynaptic transcripts. (**C**) Selected Gene Ontology (GO) annotations of transcripts significantly enriched by vGLUT1^+^ sorting. (**D**) Selected Gene Ontology (GO) annotations of transcripts significantly depleted by vGLUT1^+^ sorting. (**E**) Comparison of the percentage of mRNA containing FMR1 sites binding in the forebrain transcriptome (input) or contaminant transcriptome (depleted) or vGLUT1^+^-enriched presynaptic transcriptome (enriched). (**F**) Schematic representation of a vGLUT1^+^ presynaptic terminal with the localization of a subset of proteins coded by mRNA detected using Next Generation RNA sequencing of synaptosomes. Note that many presynaptic active zone-related mRNAs are enriched by the sorting procedure (green) whereas synaptic vesicle-related mRNAs are either significantly depleted (magenta) or not enriched by sorting. (**G**) FISH conducted on isolated vGLUT1^+^ synaptosomes validating the presence of *Rapgef4, Adcy1, Bsn, Kif5* and the absence of *Actb* mRNA in vGLUT1^+^ terminals. (**H**). Bar graph showing the analysis of FISH data indicating the percentage of vGLUT1^+^ synaptosomes that possess the indicated mRNA (*n^Rapgef4^* = 939, *n^Adcy1^* = 551, *n^Bsn^* = 1437, *n^Kif5^* = 1292 and *n^Actb^* = 359 vGLUT1^+^ terminals from at least 2 biological replicate experiments). Data are shown as mean +/− SD (Rapgef4 = 55.6% +/− 12.6; Adcy1 = 56.5% +/− 12.3; Bsn = 41.3% +/− 10.4; Kif5a = 38.4% +/− 14.7; Actb = 4.0% +/− 6.0). * Kruskal-Wallis nonparametric test followed by Dunn’s multiple comparison test, p ≤ 0.05. All scale bars = 5 μm.

## Abundant protein synthesis is detected in presynaptic terminals

To obtain direct evidence for protein synthesis in synaptic compartments, particularly presynaptic boutons, we adapted the puromycin-based metabolic labeling strategy (*33*) for detection with electron or expansion microscopy. Cultured hippocampal neurons were briefly labelled with puromycin and then fixed and processed for electron microscopy (EM) using immunogold labeling with an anti-puromycin antibody (see Methods). Using transmission EM, we were able to identify, based on morphological features (see Methods), both dendrites and synapses, including presynaptic boutons, in the images (Fig. 5A). A high fraction of presynaptic boutons and postsynaptic spines contained puromycin-positive gold particles indicating active protein synthesis within the last 10 minutes (Fig. 5A-C). The inclusion of the protein synthesis inhibitor anisomycin or the omission of puromycin, led to a dramatic reduction in the detected gold particles (Fig. 5A-C).

**Fig. 5.**
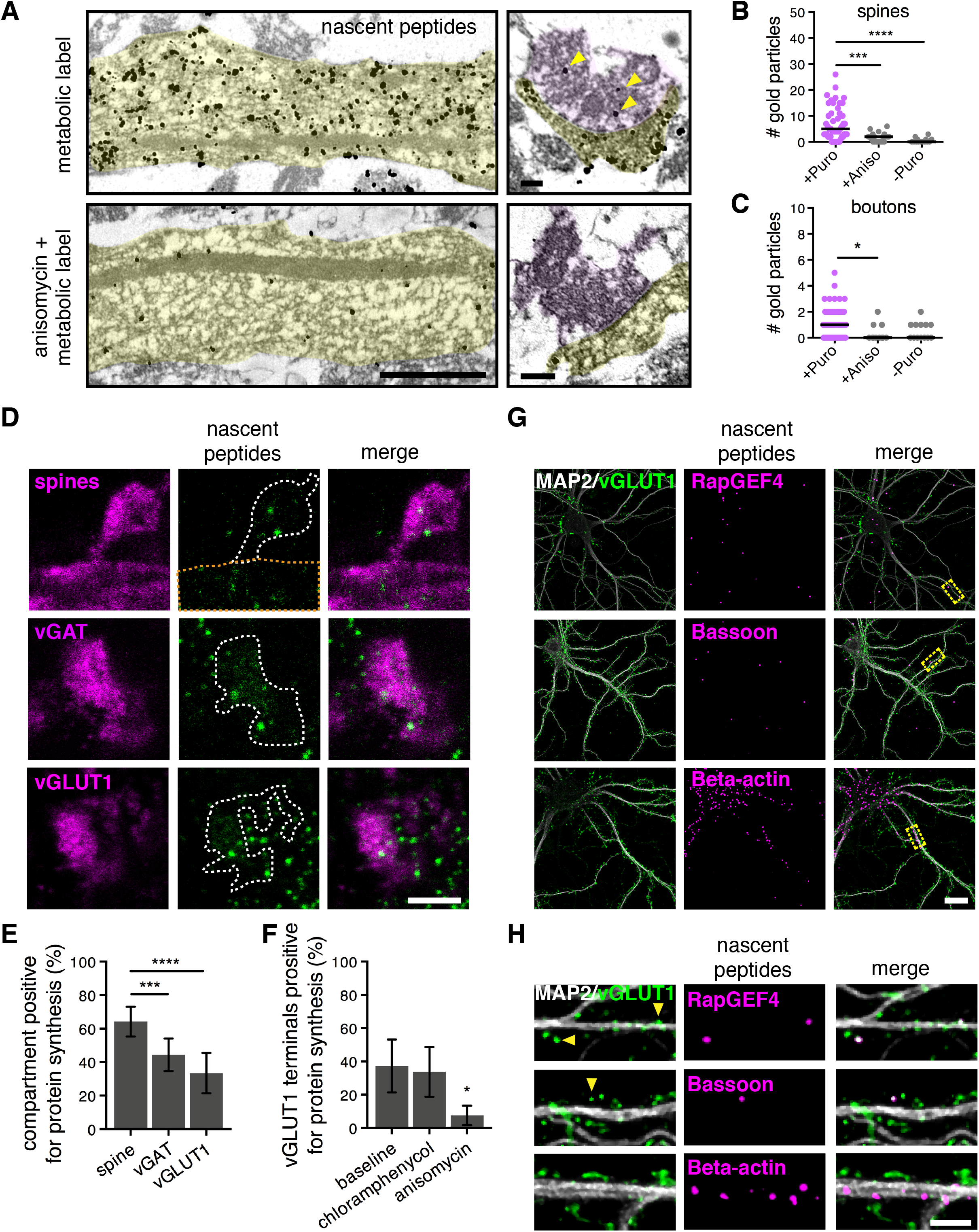
In vivo isolated pre- and postsynaptic compartments actively translate protein in the absence of external stimulation. (**A**) Electron microscope (EM) images of cultured hippocampal neurons metabolically labeled with puromycin for 10 min and then detected using immunogold (see Methods). Electron-dense particles indicate sites of protein synthesis. Shown are dendritic segments (left) and synapses (right) with gold particles present throughout the dendrite as well as in both the presynaptic (shaded lavender) and postsynaptic (shaded pale green) compartments in the absence of anisomycin. Yellow arrows indicate protein synthesis sites in presynaptic boutons. Scale bars = 1 and 0.2 μm, for dendrite and synapse, respectively. (B-C) Plots indicating the number of nascent protein immunogold labeling and the corresponding median in dendritic spines (n^+puro^ = 49, n^+aniso^ = 21, n^-puro^ = 14) (**B**) and presynaptic boutons (n^+puro^ = 54, n^+aniso^ = 11, n^-puro^ = 13) (**C**) in presence of puromycin (10 min incubation) with or without the translation inhibitor anisomycin (40 min incubation in total) and in the absence of puromycin (spines: +puro = 5, aniso = 2, -puro = 0; boutons: +puro = 1, aniso = 0, - puro = 0). Quantifications were obtained from 2 biological replicates. * Kruskal-Wallis nonparametric test followed by Dunn’s multiple comparison test, p ≤ 0.05. (**D**) Representative images of expanded cultured hippocampal neurons following 5 min of metabolic labeling and immunolabeling showing nascent protein detected in dendritic spines, excitatory presynaptic boutons or inhibitory presynaptic boutons. (**E**) Quantifications of metabolic labeling showing that a large fraction of both pre- and postsynaptic compartments were translationally active within 5 min of metabolic labeling. Data are shown as mean +/− SD (spine = 64.2 +/− 8.8, vGAT = 44.4 +/− 9.7, vGLUT1 = 33.5 +/− 12.0). ***Kruskal-Wallis nonparametric test followed by Dunn’s multiple comparison test, p ≤ 0.001. (**F**) Quantification of puromycin occupancy revealed that the inhibition of mitochondrial protein synthesis had no effect in vGLUT1+ puncta. Data are shown as mean +/− SD (baseline = 37.3 +/− 15.9, chloramphenicol = 33.7 +/− 15.0, anisomycin = 7.6 +/− 5.8). n= 3 biological replicates, 300 vGLUT1+ terminals quantified. Dunnett’s multiple comparison test, p value: *≤ 0.05. (F-G) Representative images showing newly synthesized proteins-of-interest, RapGEF4 and Bassoon, which were also identified as transcripts enriched in the vGLUT1+ transcriptome (Fig. 4B,F,G). Scale bars = 20 and 5 μm, for (**G**) and (**D**)(**H**) respectively.

The thin nature of the EM sections precludes a 3D analysis and could result in an under-estimation of ongoing protein synthesis in pre- and postsynaptic compartments. Thus, to address the frequency of translation in a well-resolved 3D volume of both presynaptic boutons and dendritic spines we used metabolic labeling in expanded cultured hippocampal neurons (Fig. 5D). Together with 5 min of metabolic labeling of nascent protein synthesis, we conducted immunocytochemical analyses using pre- (vGLUT1 or vGAT, excitatory or inhibitory) and postsynaptic (mCherry volume fill) labeling. By analyzing the coincidence of the synaptic markers with the metabolic label (again, resolved in individual z-sections), we discovered that an average of ~37 and 61% of excitatory pre- and postsynaptic compartments, respectively, and ~44% of inhibitory presynaptic terminals underwent active translation. The protein synthesis signal was dramatically reduced by the addition of the protein synthesis inhibitor anisomycin. Because mitochondria were detected in ~48% of all presynaptic terminals (*34*), we asked whether any of the presynaptic metabolic label corresponded to mitochondrial protein synthesis. The positive metabolic label in presynaptic terminals (that overlaps with either vGLUT1 or vGAT compartments) was resistant to chloramphenicol (an inhibitor of prokaryotic/mitochondrial protein synthesis) (Fig. 5F). Consistent with this, immunodetected mitochondria (anti-TOMM20 antibody) did not overlap with either the vGLUT1 or vGAT immunolabelled compartments.

We next validated the local translation of some specific candidate mRNAs, identified in the presynaptic transcriptome (Fig. 4), using Puro-PLA (*35*) together with immunolabeling to identify postsynaptic and pre-synaptic compartments (anti-Map2 and vGLUT1 antibodies, respectively). With just 5 minutes of metabolic labeling, we visualized, for example, the synthesis of both RapGEF4 and Bassoon in presynaptic compartments (Fig. 5G-H). Thus, excitatory and inhibitory presynaptic boutons (as well as postsynaptic spines) exhibited local translation with a high frequency, in the absence of any exogenous stimulation.

## Differential compartment-specific regulation of protein synthesis by plasticity

Local translation is required for several forms of synaptic plasticity including, but not limited to, potentiation induced by neurotrophins (*36*), and depression induced by metabotropic glutamate receptor activation (mGluR_1/5_) (*37*) or endocannabinoids (*16*). Capitalizing on our ability to visualize the protein synthesis that occured in three different synaptic compartments (the dendritic spine and both excitatory and inhibitory presynaptic boutons), we examined the translational signature of these three different forms of plasticity. We treated cultured hippocampal or cortical neurons with brain-derived neurotrophic factor (BDNF), an mGluR_1/5_ agonist ((S)-3,5-Dihydroxyphenylglycine hydrate; DHPG) or an endocannabinoid CB1 receptor agonist (arachidonyl-2-chloroethylamide; ACEA), adding a metabolic label for the last 5 minutes of each treatment (Fig. 6A). Immunocytochemical detection of nascent protein and markers of each synaptic compartment was conducted and the samples were then subjected to expansion microscopy. The pattern of protein synthesis in the three compartments of interest, in both brain areas, indicated that each type of plasticity yielded a unique constellation of synaptic translation loci: BDNF caused an increase in local translation in dendritic spines and both excitatory and inhibitory boutons (Fig. 6B-E), DHPG caused an increase in dendritic spines only, and ACEA caused an increase in inhibitory boutons exclusively (Fig. 6C-E). The addition of anisomycin significantly reduced the signal in all conditions. This pattern, while clearly evident in the synaptic compartments, was not observed in either the soma or the total dendrite, strongly suggesting a synaptic, localized response. In separate experiments, we examined whether the above three agonists changed the distribution or amount of poly(A) mRNA and found no significant change. Thus the compartment-specific translation observed above was mediated by local, enhanced translation of mRNAs already resident at the synapse. Furthermore, our results provide subcellular resolution on the local proteomic remodeling that drives different forms of synaptic plasticity.

**Fig. 6.**
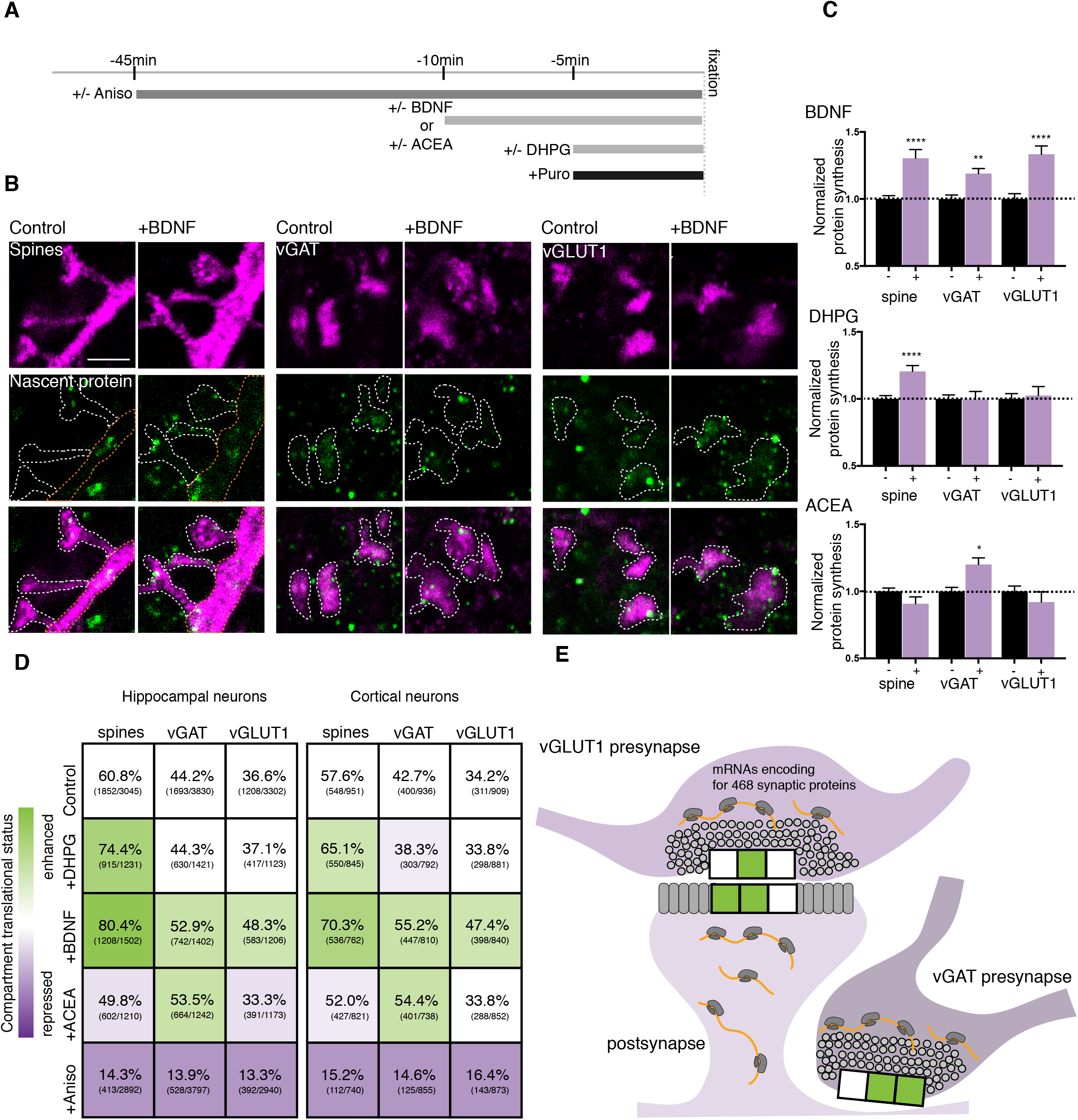
Compartment-specific translation patterns decode different forms of plasticity. (**A**) Scheme showing the timing of the different plasticity induction protocols and the metabolic labeling (+puro). (**B**) Representative images from hippocampal cultures showing both the immunostained, and metabolically labelled compartments indicating newly synthesized protein following expansion microscopy for one of the plasticity conditions (+/− BDNF). Scale bar = 5 μm. (**C**) Bar graphs indicating the specific translation pattern in different subcellular compartments (vGAT^+^ or vGLUT1^+^ presynaptic terminals or spine) following the 3 different plasticity treatments, normalized to the control condition in hippocampal cultures, n = 4-6 biological replicates per condition, for total puncta counted see numbers in the matrix shown in D. Unpaired t-tests, p values: ** ≤ 0.01, *** p ≤ 0.0001. (**D**) Matrices for hippocampal and cortical neurons showing both the synaptic compartment (spine, excitatory presynaptic compartment or inhibitory presynaptic compartment) and the plasticity agonist (BDNF, DHPG or ACEA) applied and the percentage of compartments that exhibited protein synthesis. Shown in parentheses are the numbers of labeled compartments over the total number of compartments examined. Colors represent the change in protein synthesis-with green and lavender colors indicating a stimulation or repression of protein synthesis, respectively (see color look-up table). (**E**) Summary scheme indicating how each different form of plasticity examined has a specific translational signature. The 3 compartments represented by the horizontal boxes indicate the stimulation of protein synthesis by DHPG, BDNF, or ACEA, in that order.

## Discussion

Here, we investigated the localization and stimulation of protein synthesis in mature synapses and unambiguously identified protein synthesis machinery and translation in individual presynaptic compartments from 3 different brain areas. In adult rodent brain slices and cultured hippocampal neurons we found that over 75% of both excitatory and inhibitory presynaptic terminals (see also (*16*) contained rRNA, ribosomes and poly(A) mRNA. Both light (confocal and super-resolution) and electron microscopy revealed in the absence of overt stimulation a surprisingly high level of ongoing protein synthesis in both pre- and postsynaptic compartments: with only 5 min of labeling ~40% of both excitatory and inhibitory presynaptic terminals and ~60% of dendritic spines exhibited active translation. Puromycin, a tRNA mimic, was used to metabolically label nascent proteins (*33*); we used an optimized low concentration in order to label nascent peptides while avoiding a complete block of protein synthesis. The stringency of our presynaptic translation measurements (e.g. the requirement that the metabolic label spatially overlap with a vesicular marker immunolabeling) may also lead to an underestimate of actively translating compartments, particularly when compared to the spine measurements where a volume-filling label was used. As such, we believe that the above values likely represent a conservative estimate of the fraction of compartments undergoing translation in the labeling window. Indeed, we could increase the number of active compartments using chemical stimulations. This confirms that we have identified a “lower-bound”: we detected an even higher fraction of actively translating compartments (~81, 54 and 48 percent of spines, vGAT and vGLUT1 terminals) after 5 min of metabolic labeling.

Thousands of mRNA transcripts are present distally in neuronal processes, where they can be locally used for protein synthesis (*38–41*). Notably, the transcriptome of retinal ganglion cell axons has been characterized both during development (*42*, *43*) and the retinal ganglion cell translatome has been identified in the adult mouse (*15*). Here, using mature mouse forebrain synaptosomes that are enriched for vGLUT1^+^ presynaptic terminals we identified ~450 transcripts that were enriched, relative to the “pre-sorted”/bulk synaptosome transcriptome. There were also many transcripts shared between the pre-sorted and sorted synaptosomes that were not enrichment in the vGLUT1^+^ transcriptome but likely represent important translation targets within post- and/or pre-synaptic compartments. Within this vGLUT1^+^-enriched transcriptome, we detected many mRNAs that code for proteins that regulate vesicle release probability including *Rims, Adcy1*, and *Bsn*. Using Puro-PLA (*35*) we validated the synthesis of several presynaptic proteins, including RapGEF4 and Bassoon, in identified nerve terminals within minutes of metabolic labeling. Some of the earliest studies suggesting translation in axons observed radioactive labeling in synaptosomes (*44*, *45*). Interestingly, mRNAs coding for synaptic vesicle proteins were lacking in our vGLUT1^+^ transcriptome. Perhaps local translation of presynaptic proteins could work in concert with the well-documented transport of presynaptic proteins and complexes within axons (*46*) to supply and regulate neurotransmitter release and homeostasis in mature, healthy nerve terminals.

We detected an enrichment of transcripts in several functional categories. For example, we noted an abundance of mRNAs coding for proteins that directly regulate translation, including eukaryotic initiation and elongation factors (see also (*15*)). While many of these proteins have been detected within dendrites (*47*, *48*) whether they are present in excess or limited quantities is unknown. Signaling events at the synapse could thus boost translational capacity by synthesizing these potentially rate-limiting regulatory elements. Importantly, local protein synthesis is dysregulated in many neurodevelopmental disorders (*49*) and recent attention has focused on a presynaptic locus of some important proteins like FMRP (*50*). In this regard, we note that over 10% of the vGLUT1^+^-enriched presynaptic transcripts possess an FMRP-binding site (*32*). Shigeoka et al., also observed an abundance of FMRP targets in the retinal ganglion cell axonal translatome (*15*).

Multiple forms of synaptic plasticity involve local translation in dendrites including, BDNF-induced synaptic potentiation (*36*), mGluR-dependent long-term depression (*51*), dopamine-induced plasticity (*52*) and homeostatic plasticity (*48*) and activation of presynaptic CB1 receptors by retrograde endocannabinoid signaling stimulates local protein synthesis in inhibitory terminals to produce long-term depression of inhibitory transmission (*16*). We found local translation in both the pre- and postsynaptic compartments to be differentially regulated by 3 of the above forms of plasticity in a compartment-specific manner. These data indicate that there is also information about the recent synaptic history and the expression of plasticity in the particular pattern of translation loci in synaptic compartments. Together with the selection of particular mRNAs for translation, owing to unique regulatory elements present in the 3’UTRs (*53*), a unique and remodeled synaptic proteome for each kind of plasticity can be achieved. Our findings demonstrate that local protein synthesis is a ubiquitous feature of both sides of the synapse-it occurs in both excitatory and inhibitory presynaptic boutons, as well as dendritic spines, under basal conditions and is differentially recruited in these compartments to modify local proteomes. Taken together with the well-documented system of microtubule-based transport (in both axons and dendrites) to supply both mRNA and protein, local synthesis adds an important source of protein that presumably can be exploited to alter the local proteome with spatial and temporal precision.

## Materials and Methods

### Cultured neurons

Dissociated rat hippocampal or cortical neuron cultures were prepared and maintained as described previously (*54*). Briefly, we dissected hippocampi or cortices from postnatal day 0-1 rat pups of either sex (Sprague-Dawley strain; Charles River Laboratories), dissociated them with papain (Sigma) and plated them at a density of 40×10^3^ cells/cm^2^ on poly-D-lysine coated glass-bottom Petri dishes (MatTek). Neurons were maintained and matured in a humidified atmosphere at 37°C and 5% CO_2_ in growth medium (Neurobasal-A supplemented with B27 and GlutaMAX-I, life technologies) for 18-21 days in vitro (DIV) to ensure synapse maturation. All experiments complied with national animal care guidelines and the guidelines issued by the Max Planck Society, and were approved by local authorities. For transfection, DIV7-11 neurons were transfected with mCherry-C1 or EGFP-C1 using Effectene (Qiagen), as previously described (*55*). Transfected cells were maintained until DIV18-21 for experiments.

### Preparation of mouse brain sections

12-week-old mice were perfused with 1x phosphate buffered saline (PBS) and 4% (v/v) paraformaldehyde solution in PBS. Brains were dissected, and sliced to 2 mm and fixed for 3 h at room temperature. Slices were cryoprotected in 20% (w/v) sucrose in PBS (DEPC-treated) overnight at 4°C and cryosectioned at 20 μm thickness. Samples were then stored at −20°C in 80% ethanol until use.

### In situ hybridization in synaptosomes and cultured neurons

All steps were performed at room temperature, unless stated otherwise. Bottom glass dishes with attached neurons (DIV 21+) or pre-sorted or vGLUT1+ sorted synaptosomes plated on gelatinized coverslips were fixed in paraformaldehyde 4% in lysine phosphate buffer pH7.4 containing 2.5% sucrose for 15-20 min. Target specific in situ hybridization was performed using Stellaris™ probes (LGC Bioresearch) as previously described (*56*) and oligo(dT) and d(A) performed as in (*57*). Following fixation, cells were washed in PBS + 5 mM MgCl_2_, followed by dehydration in 80% ethanol overnight at −20°C. Samples were rehydrated in PBS + MgCl_2_, followed by 2x 1X saline-sodium citrate (SSC) washes, followed by a 5 min wash in 2X SSC + 30% formamide for 5 min. Biotin labeled probes for 18S and 28S rRNA (Stellaris, Biosearch technology), and 18mer oligo(dT)/oligo(dA) (Eurofins) were diluted into 100μl hybridization buffer and incubated on cells overnight at 37°C. Following probe hybridization, samples were washed twice in 2X SSC + 30% formamide for 30 minutes each, followed by 5x 1X SSC washes. After completion of in situ hybridization, samples were washed with PBS and subsequently processed for immunofluorescence. For RNaseA/T1 controls 1mL of digestion buffer (10 mM Tris-HCl, 300 mM NaCl, 5 mM EDTA) was added to sample with and without 20 uL of RNaseA/T1 (Thermo Fisher) for 30 min at 37°C following sample rehydration. For permeabilization controls, EtOH dehydration series were omitted.

### In situ hybridization in tissue

Following rehydration, samples were postfixed for 5 min in ice cold 4% PFA, followed by washes in 2xSSC. Samples were treated with 0.1 M triethanolamine-HCl pH 8.0 + acetic anhydride for 10 min to reduce nonspecific hybridization. Following washing in ice cold H_2_0 samples were incubated in ice cold methanol/acetone, and then washed in ice cold 1x SSC. Samples were blocked for endogenous biotin using streptavidin for 30 min at 37°C followed by a biotin was for 5 min (Thermo Fisher). Samples were then incubated in 2X SSC for 10 min, followed by 2X SSC + 50% Formamide for 1 h. FISH probes were diluted to 2X in 200 μL hybridization buffer and incubated overnight at 37°C. Samples were washed 5x in 2X SSC + 50% formamide for 60 min each at 37°C, followed by 5 washes in 2X SSC for 10 min each. After completion of in situ hybridization, samples were washed with PBS and subsequently processed for immunofluorescence.

### Immunofluorescence in synaptosomes and cultured neurons

All steps were performed at room temperature, unless stated otherwise. Bottom glass dishes with attached neurons (DIV 18-21) or pre-sorted or vGLUT1+ sorted synaptosomes plated on gelatinize coverslips were fixed in paraformaldehyde 4% in lysine phosphate buffer pH7.4 containing 2.5% of sucrose for 15-20 min. Cells were then permeabilized for 10 min in PBS + 0.5% Triton-X 100 (Sigma). Samples were incubated in blocking buffer (4% goat serum in PBS) or biotin-free blocking buffer (4% biotin free BSA in PBS, for FISH experiments) for 30 min. After 3 washes in PBS for 5 min each, samples were incubated in blocking buffer (4% goat serum in PBS for cell culture experiments, 4% biotin free BSA in PBS for cell culture FISH experiments) for 1 to 2 h with secondary antibodies. We used the following antibodies: guinea pig anti-MAP2 (Synaptic Systems, 1:2000), rabbit anti-biotin (Bethyl, 1:1000), rabbit anti-biotin (Cell Signaling, 1:1000), chicken anti-GFP (Aves, 1:1000), chicken anti-mCherry (Abcam, 1:1000), mouse anti-PSD-95 (Thermo Fisher Scientifics, 1:1000), mouse anti-synaptopodin (Merck, 1:500), rabbit anti-calreticulin (Abcam, 1:1000), guinea pig anti-Homer1 (Synaptic Systems, 1:1000), mouse anti-Puromycin (Kerafast, 1:500-1000), guinea pig anti-VGLUT1 (Synaptic Systems, 1:500-2000), guinea pig anti-VGAT (Synaptic Systems, 1:500-2000), rabbit anti-VGAT (Synaptic Systems, 1:1000), mouse anti-Smi-312 (Covance, 1:2000), rabbit anti-RPS11 (Bethyl, 1:200), rabbit anti-RPL26 (Sigma, 1:500-1000). After 3 washes in PBS for 5 min each, samples were incubated in blocking buffer (4% goat serum in PBS) for 1 to 3 h at room temperature with secondary antibodies. For expansion microscopy, we used the following dyes coupled to our secondary antibodies: Alexa-488, Alexa-568, Abberior STAR-635.

### Immunofluorescence in tissue sections

Brain sections were incubated in 4% goat serum + 0.5% Triton-X 100 for normal IF or 4% biotin free BSA + 0.5% Triton-X 100 for FISH experiments at room temperature for 4 h. Primary antibody staining was carried out overnight in the same buffer at 4°C. Samples were washed 5x in PBS before carrying out secondary antibody staining for 3 h at room temperature.

### Total protein labeling

Prior to permeabilization cells or tissue were incubated with 0.2 M bicarbonate buffer supplemented with 0.5 mg/mL Alexa568 NHS Ester (Thermo) for 15 min at room temperature, to label all amine groups in the sample with the Alexa dye. Samples were washed 5x with PBS and used for subsequent IF or FISH experiments.

### Cell treatments

For puromycin labeling experiments, cultured neurons were treated with 10 μM puromycin for 5 min, if not stated otherwise. Anisomycin treatment (40 μM) was performed 20-45 min prior to puromycin labeling. BDNF (50 ng/mL) was added for 10 min, arachidonyl-2-chloroethylamide (ACEA; 50 μM) was added for 10 min and (S)-3,5-Dihydroxyphenylglycine hydrate (DHPG; 50 μM) was added for 5 min. For mitochondrial protein synthesis inhibition experiments, 40 μM chloramphenicol was added for 40 min prior to the addition of puromycin.

### Expansion microscopy

Following immunofluorescence labelling, samples were treated with Acryloyl-X-SE (Thermo Fisher) overnight at room temperature. Following washing steps, 200μl of monomer solution was added to the coverslip and gelation was carried out at 37°C for 1h. For tissue sections, water was replaced with 4-hydroxy TEMPO (Thermo Fisher) as previously described (*22*). Tissue sections were pre-incubated in monomer solution at 4°C for 30 min, prior to transferring the samples to 37°C for 2 h to allow gelation to occur. Following ProteinaseK (NEB) digestion overnight, slightly expanded gels were transferred to a larger dish and water exchange was performed until gels were fully expanded. Expanded gels were transferred into 50×7 mm glass bottom dishes (WillCo Wells) for imaging. Expanded gels were imaged using Zeiss LSM780/880 confocal microscopes and a 63x oil objective (NA 1.4, PSF: LSM780-0.240/0.258/0.729 μm; LSM880-0.252/0.203/0.563 μm x/y/z) for cultured cells and synaptosomes and a 40x oil objective (NA 1.3, PSF: LSM780-0.217/0.260/0.566 μm; LSM880-0.238/0.253/0.636 μm x/y/z). Z-stacks (0.37 μm 63x/ 0.43 μm 40x) spanning the entire volume of imaged neurons, synaptosomes or tissue were obtained and analyzed using Imaris (Bitplane) and ImageJ.

### Image analysis

To assess signal occupancy in spines or boutons, the compartment was considered positive for either puromycin or RNA if signal was detected in at least 3 individual consecutive z slices. Presynaptic terminals were defined by vGLUT1 and vGAT signal, and spines were defined based on morphology from mCherry or GFP volume filling. To be considered a spine, the compartment must be a clearly defined protrusion from the dendrite-extending at least 1.5um away (in the expanded images) from the dendritic shaft. For tissue sections, due to the increased density and complexity of the samples, images were first processed in Imaris. 3D surface masks were generated corresponding to either vGLUT1 or vGAT signal. These presynaptic surface masks were used to generate a new channel, corresponding to the puromycin or RNA signal found within the 3D presynaptic volume. These images were then compressed into max intensity projections, and the number of positive vGLUT or vGAT terminals were scored, positive or negative, based on the presynaptic puromycin or RNA channel. To assess the amount of signal falling within and outside of cells, samples with total protein labeled (Alexa-568 NHS ester-see total protein labeling section) and FISH were analyzed in Imaris. The total protein Alexa-568 channel was used to make a 3D surface mask, and all RNA FISH signal falling within this mask was copied into a new third channel. The total signal intensity of the original RNA FISH signal channel was then measured as well as the signal corresponding to the FISH signal within cells. The ratio of cellular signal/total signal was used to assess the fraction of signal falling within cells. For soma and dendrite relative puromycin incorporation measurements, sum intensity projections were made and using the mCherry signal as a mask. The soma or the dendrite (starting 15 μm away from the cell body, and extending for at least 70 μm) was selected and total puro signal was assessed and normalized to the dendritic or soma area.

### Proximity ligation assay (PLA)

Detection of newly synthesized proteins by proximity ligation was carried out using anti-puromycin antibodies (mouse anti-Puromycin from Kerafast, 1:500-1000) in combination with protein-specific antibodies (rabbit anti-RapGEF4 from Invitrogen, 1:250; rabbit anti-bassoon from Enzo, 1:500; rabbit anti-beta-actin from Abcam, 1:1000). We used Duolink reagents (Sigma) and followed the protocol provide by the manufacturer with some modifications described below. We routinely used rabbit PLA^plus^ and mouse PLA^minus^ probes as secondary antibodies and the “Duolink Detection reagents Red” (Sigma) for ligation, amplification and label probe binding. Briefly, after a brief 5min of metabolic labeling, hippocampal cultured neurons (21 DIV) were fixed in PBS-sucrose, permeabilized in PBS + 0.5% Triton-X 100 and blocked in PBS + 4% goat serum as described previously for immunocytochemistry assays. Then, neurons were incubated overnight at 4°C in PBS + 4% goat serum containing primary antibodies: mouse anti-puromycin, rabbit anti-protein of interest (e.i. anti-RapGEF4, Bassoon or beta-actin), chicken anti-MAP2 (Abcam, 1:2000) and guinea-pig anti-vGLUT1 (Synaptic System, 1:1000). After washing, PLA probes were applied in 1:10 dilution in PBS with 4% goat serum for 1 h at 37 °C, washed several times with wash buffer A (0.01 M Tris, 0.15 M NaCl, 0.05% Tween 20) and incubated for 30 min with the ligation reaction containing the circularization oligos and T4 ligase prepared according to the manufacturer’s recommendations (Duolink Detection reagents Red, Sigma) in a prewarmed humidified chamber at 37°C. Amplification and label probe binding was performed after further washes with wash buffer A with the amplification reaction mixture containing Phi29 polymerase and the fluorophore-labeled detection oligo prepared according to the manufacturer’s recommendations (Duolink Detection reagents Red, Sigma) in a prewarmed humidified chamber at 37 °C for 100 min. Amplification was stopped by three washes in wash buffer B (0.2 M Tris, 0.1 M NaCl, pH 7.5). We note that for better signal stability, cells were kept in wash buffer B at 4°C until imaging.

### Pre-embedding immuno-detection of newly synthesized proteins visualized by electron microscopy

For the detection of newly synthetized proteins in neurons, we performed pre-embedding immuno-detection of puromycin as described below. All steps were performed at room temperature, if not stated otherwise. Bottom glass dishes with attached neurons (DIV 28) were fixed in paraformaldehyde 4% and 0.05% glutaraldehyde in 0.2 mM HEPES buffer pH7.2 for 45 min. Cells were then permeabilized for 10 min in PBS containing 0.5% Triton-X 100 (Sigma). Fixation reagents were quenched using freshly made 1 mg/ml borohydride in HEPES 0.2 mM pH8 for 10 min. Antibodies were applied on the samples in blocking buffer (PBS + 2% biotin-free BSA). After 30 min in blocking buffer cells were incubated with mouse anti-puromycin (Kerafast, 1:2000) for 1 h at room temperature. Prior to the 1 h incubation at room temperature with anti-mouse antibody coupled to biotin (Abcam, 1:1000) we performed an endogenous biotin-block (Thermo Fisher). Biotin was detected with a rabbit anti-biotin antibody coupled to 1 nm nanogold particles (1:100, FluoroNanogold Alexa-594, Nanoprobes). Samples were post-fixed in 1% glutaraldehyde in 0.2 mM HEPES pH = 7.2 for 30 min and fixation was quenched with 100 mM Glycine in PBS for 10 min. Samples were then washed in water three times and then three times with 20 mM sodium citrate buffer pH = 7.0. Nanogold particles were subsequently amplified using silver amplification for 6 min (Serva) and fixed again in 0.2% OsO4 for 30 min. Samples were then stained with 0.25% uranyl acetate (Serva) in the dark for 30 min. After washing and dehydrating with ethanol, samples were embedded in Epon (Serva). Sections (60 nm thick) were mounted onto Formvar-coated copper grids (Serva). Grids were imaged with a transmission electron microscope LEO (Zeiss) 912 OMEGA.

### Synaptosome isolation

Synaptosomes were generated from 6-8 weeks old mouse forebrain, hippocampus or cerebellum of wild-type and VGLUT1^VENUS^ knock-in mice as described previously (*27*, *28*, *30*). Our synaptosome preparation was chosen and adapted from our previously published protocol (*27–29*) to favor the isolation of synaptosomes with presynaptic compartments with closed membrane bilayers and postsynaptic compartments with open membrane bilayers. Briefly, the cerebellum, forebrain, or both hippocampi from a single mouse was homogenized in 2 mL of ice-cold homogenization buffer (0.32 M sucrose, 4 mM HEPES pH 7.4, protease inhibitor cocktail EGTA-free (Calbiochem, 1:1000), RNasin (Promega, 1:1000) using a 2 mL glass-Teflon homogenizer with 12 gentle strokes. The homogenizer was then rinsed with additional 3 mL of homogenization buffer, and the combined 5 mL of homogenate was centrifuged at 1000 g for 8 min at 4°C. The supernatant was centrifuged again at 12,500 g for 15 min at 4°C. The synaptosome-enriched pellet was then re-suspended in 1 mL of homogenization buffer. This fraction was finally layered on top of a two-step sucrose density gradient (5 mL of 1.2 M and 5 mL of 0.8 M sucrose, HEPES 4 mM, protease inhibitor cocktail EGTA-free, as above). The gradient was centrifuged at 50,000 g for 70 min at 4°C. Synaptosomes were recovered through the tube wall, at the interface of 0.8 and 1.2 M sucrose, using a syringe to minimize contamination with lighter fractions enriched in myelin. The resulting fraction is referred as pre-sorted (sucrose) synaptosomes (or S-synaptosomes) as opposed to vGLUT1^+^ sorted synaptosomes (or FASS-synaptosomes).

### Post-embedding immuno-detection of ribosomal protein RPS11 visualized by electron microscopy

Pre-sorted synaptosomes (~750 μl) on iced were mixed with ice cold PBS containing protease inhibitor cocktail EGTA-free (Calbiochem, 1:1000) and RNasin (Promega, 1:1000) to obtain a final volume of 1.5 ml. Pre-sorted synaptomes were centrifuged at 16.8 g for 5 min at 4°C. All the subsequent steps were performed at room temperature unless stated otherwise. The resulting pellet was then fixed using in PBS + 4% PFA + 2% glutaraldehyde for 1 h. The pellet was then cut in 2 to 4 pieces, that were then washed 4x in PBS and 5x in H2O before post-fixation in 0.2% OsO4 in water for 30 min. Samples were then stained with 0.25% uranyl acetate (Serva) in the dark for 30 min. After washing and dehydrating with ethanol, samples were embedded in Epon (Serva). Sections (60 nm thick) were mounted onto nickel grids (Serva) and processed for immunostaining. After 3 washes in TBST, grids were blocked for 10 min in TBST with 10% Normal Goat Serum (NGS). Antibodies were applied in TBST with 1% NGS. Grids were incubated with anti-RPS11 (Bethyl, 1:200) overnight in the dark. The next day grids were washed 5x 3 min with TBS before being incubated with anti-rabbit antibody coupled to ~10mn colloidal gold (BBI solution, 1:50) for 2 h. Finally, grids were washed 3x 5min in TBS, 3x 5min PBS, 3x 5 min in H_2_O and further stained with 0.4% uranyl acetate (Serva) and contrasted with lead citrate (Merck). Grids were imaged with a transmission electron microscope LEO (Zeiss) 912 OMEGA. Control grids in which the primary antibody against RPS11 was omitted yielded no gold signal in presynaptic terminals.

### Fluorescence-activated synaptosomes sorting (FASS)

S-synaptosome sorting was performed as described previously (*27*, *28*). The FACSAria-II (BD Biosciences) operated using a 70 μm nozzle. Briefly, S-synaptosomes were stored on ice and diluted in PBS containing protease and RNAase inhibitor as above and labeled with a red (excitation/emission maxima ~515/640 nm) lipophilic dye FM4-64 (Thermo Fisher Scientific, 1.5 μg/mL). Dilution was optimized to obtain an event rate of 20,000 – 25,000 events/s. FM4-64 was used to trigger the FACSAria detection on all biological membranes in the sample. A first gate delineated small particles (singlets) and excluded events showing correlated high values for FSC and SSC area (aggregates and large particles). The “singlets” gate was sub-gated according to vGLUT1^VENUS^ fluorescence intensity using the 488 laser line. Thus “singlets” were sorted into two fractions, the vGLUT1^VENUS^ negative fraction (vGLUT1^-^) and the vGLUT1^VENUS^ positive fraction (vGLUT1^+^). Those two fractions were subsequently either collected onto filters and processed for RNA Next Generation Sequencing (NGS) or plated onto gelatinized coverslips at a density of 1 Mio particles per 12 mm coverslips by centrifugation at 6,800 g for 34 min in 24 well plates and then processed for immunofluorescence and/or in situ hybridization (see protocol above).

### Stimulation Emission Depletion microscopy (STED)

Super resolved images of vGLUT1^+^-sorted synaptosomes were obtained using a Leica SP8 WLL2 inverted DMI6000 confocal microscope (Leica Microsystems, Mannheim, Germany) equipped with the 3D STED module. In the STED module, we used a 775 nm laser line to deplete Alexa-594 and ATTO-647N. We achieved two color STED with a final ~40 nm resolution using a 93X glycerol objective NA 1.30 and a white light laser 2 (WLL2) with freely tunable excitation from 470 to 670 nm (1 nm steps) and a diode laser at 405 nm. The microscope was equipped with 2 internal PMTs and 2 internal hybrid detectors.

### RNA Next Generation Sequencing

Total RNA were extracted using TRIzol LS reagent (Thermo Fisher) and Direct-zol RNA microprep kit (Zymo) for four different samples: input (mouse forebrain) from wildtype and VGLUT1^VENUS^ mice, pre-sorted synaptosomes from wildtype and VGLUT1^VENUS^ mice; and vGLUT1^+^ sorted synaptosomes VGLUT1^VENUS^ mice P3 fractions (after filtration to remove the excess of PBS). Total RNA libraries were generated using NEBNext rRNA Depletion kit combined with NEBNext Ultra II Direction RNA Library Prep kit for Illumina (New England Biolabs). We used 100 ng of starting RNA material for input and pre-sorted synaptosomes and 1-5 ng for vGLUT1^+^ sorted synaptosomes which corresponds to ~100 Mio synaptosomes collected per P3 fractions. In the final amplification step of the library preparation or PCR enrichment step we used 12 cycles of amplification for the input and pre-sorted synaptosomes and 16 cycles of amplification for amplification the vGLUT1^+^ sorted synaptosomes. Ultimately we obtained cDNA libraries of ~250 bp with each sample containing a specific barcode. Libraries corresponding to replicates 1 and 2 were sequenced together in the same sequencing run. The libraries corresponding to replicates 3 and 4 (only used as an input sample) were sequenced in a subsequent run. For sequencing we used 10 ng of starting material for each library in a high throughput Illumina flow cell using Illumina NextSeq 550 Instrument.

### RNA Sequencing data analysis

For detection and annotation of the sequencing reads we used the following pipeline:

i. *Genome alignment*. Reference genome was mouse version mm10 from UCSC (link: http://hgdownload.soe.ucsc.edu/goldenPath/mm10/bigZips/). Reads alignment was conducted with STAR aligner (*58*) (version 2.5.2) with the following parameters: STAR --runMode alignRead --genomeDir $path_genome_index_mm10 -- readFilesCommand zcat --outStd Log --outSAMtype BAM SortedByCoordinate -- outSAMstrandField intronMotif --outFilterIntronMotifs RemoveNoncanonical -- alignSoftClipAtReferenceEnds No --outFilterScoreMinOverLread 0.25 -- outFilterMatchNminOverLread 0.25
ii. *Annotation assignment*. Annotation GTF file was downloaded from UCSC Table Browser Tool with the following parameters: Clade: Mammal, genome: Mouse, assembly: Dec. 2011 (GRCm38/mm10), group: Genes and Gene Predictions, track: NCBI RefSeq, table: RefSeq All (ncbiRefSeq), output format: GTF – gene transfer format (limited) (link: http://genome.ucsc.edu/cgi-bin/hgTables?hgsid=706713641_R7J6gGZbwmO18wecxAAiXglgAlzN&clade=mammal&org=Mouse&db=mm10&hgta_group=genes&hgta_track=refSeqComposite&hgta_table=0&hgta_regionType=genome&position=chr12%3A56694976-56714605&hgta_outputType=primaryTable&hgta_outFileName=) Gene expression was counted using featureCounts (link: http://bioinf.wehi.edu.au/featureCounts/) parameters: featureCounts -a path_genome_annotation -o counts.txt -t exon -Q 255 -T 12 $(ls /path/bams/files/*.bam)
iii. *Differential Expression analysis*. Differential Expression analysis was performed using DESeq2 in R (available at https://bioconductor.org/packages/release/bioc/html/DESeq2.html) setting the differential expression cut-off to 1.3-fold change and false discovery rate q ≤ 0.1 using the Benjamini-Hochberg method (*59*, *60*).
iv. *Detection of poly(A/T) repeats*. A fuzzy polyA match algorithm was adapted from Kent WJ, GenomeRes. 2002 (https://www.ncbi.nlm.nih.gov/pubmed/11932250), Iterator will start at every position on the sequence with two consecutive Adenine (A) (or thymine (T) respectively), setting initial score to 10. Every match adds a score of 1 and a mismatch adds a score of −8. The iterator will stop incrementing when the score value drops below 0. The longest span of the iterator is kept per transcript as a measure of detected repeat sequence.

All Venn diagrams were obtained using Venn 2.1 web tools (*61*).

## Acknowledgements

We thank I. Bartnik, N. Fuerst, A. Staab, D. Vogel and C. Thum for the preparation of cultured neurons, S. tom Dieck and T.W. Lee for important work in preliminary studies, Ha Nguyen for experiments shown in Figure 3J, Maria Florencia Angelo, Anaëlle Stum and Vincent Pitard for technical assistance with FASS experiments (Flow cytometry facility, CNRS UMS 3427, INSERM US 005, Univ. Bordeaux, F-33000 Bordeaux, France), Fabrice Cordelières for image analysis routines (Bordeaux Imaging Center, Univ. Bordeaux, F-33000 Bordeaux, France) and Georgi Tushev for help with RNA seq analyses.

## Author contributions

A.S.H. and P.D.A. designed, conducted and analyzed experiments. E.H. and B.L. designed and supervised experiments. E.M.S. designed experiments, supervised the project, and wrote the paper. All authors edited the paper.

